# T2R5 agonist phendione decreases cell viability and induces apoptosis in head and neck squamous cell carcinoma

**DOI:** 10.1101/2025.05.14.654120

**Authors:** Sarah Sywanycz, Brianna L. Hill, Zoey A. Miller, Gavin Turner, Lily Huang, Kyle Polen, Robert J. Lee, Ryan M. Carey

## Abstract

Bitter taste receptors (T2Rs), a family of G-protein coupled receptors, are emerging as potential therapeutic targets in head and neck squamous cell carcinoma (HNSCC). Phendione, a known T2R5 agonist, has not been previously investigated in HNSCC. Here, we show that phendione activates endogenously expressed T2R5 in HNSCC cells and *ex vivo* tumor samples, inducing sustained calcium responses, reducing cell viability, and promoting apoptosis through a T2R5-dependent mechanism. Analysis of The Cancer Genome Atlas data revealed that high T2R5 expression in HNSCC tumors correlates with improved long-term disease-specific survival, suggesting a potential tumor-suppressive role for T2R5. These findings highlight T2R5 as a promising therapeutic target in HNSCC and support further investigation of phendione or other T2R5 agonists as potential anti-cancer agents.

## Introduction

Head and neck squamous cell carcinomas (HNSCCs) are a group of malignancies arising from the epithelial tissues of the oral cavity, pharynx, larynx, and sinonasal cavity (1). HNSCC ranks as the seventh most common cancer globally, with a five-year average survival of 60-70%. Major risk factors for HNSCC include tobacco and alcohol use, as well as human papilloma virus (HPV) (1,2). Despite public health efforts, such as HPV vaccination and smoking cessation initiatives (3,4), the incidence of HNSCC is projected to rise by approximately 30% annually by 2030 (2,5). Current treatments, including surgery, chemotherapy, and radiation therapy, are often associated with significant long-term impacts on patients’ quality of life (6), underscoring the need for novel, less invasive therapeutic strategies.

Bitter taste receptors (taste family 2 receptors or T2Rs) are a family of G-protein coupled receptors (GPCRs) with 26 functional isoforms (7,8). While T2Rs are traditionally associated with detecting bitter compounds on the tongue, they are also expressed in extra-oral tissues (9–11) where they may regulate diverse physiological and pathological processes. Recently, we and others have shown that T2Rs regulate cell proliferation in malignancies such as HNSCC (10,12–15). T2Rs frequently induce calcium (Ca^2+^) release from endoplasmic reticulum (ER) stores via inositol-1,4,5- triphosphate (IP3) (16,17). In HNSCC, T2R isoform 14 and 4 (T2R14 and T2R4) activation induce apoptosis through mitochondrial Ca^2+^ overload, reactive oxygen species (ROS) production, and caspase-3/7 activation (13,14). Notably, preclinical evidence suggests that T2R14 agonist, lidocaine, may activate apoptosis in cancer cells and/or inhibit metastasis (18–20), prompting clinical trials to evaluate its efficacy in solid tumors including breast cancer (21) and HNSCC (22).

Unlike T2R14, which is activated by over 100 known bitter compounds (23,24), other T2R isoforms demonstrate greater ligand selectivity. T2R5, encoded by the *TAS2R5* gene has only 12 identified agonists, including heterocyclic metal chelator 1,10-phenanthroline (25–28). Among T2R5 agonists, the derivative 1,10-phenanthroline-5,6-dione (phendione) exhibits relatively high potency for T2R5 in human airway smooth muscle cells (29). Phendione-based therapies have also been studied as effective antibacterial, antifungal, and antitumor agents (30–34). In contrast to cisplatin, a commonly used chemotherapeutic agent that is first line for HNSCC, phendione does not cause double stranded DNA-breaks (35–37). Instead, phendione exerts its potential antitumor effects through a distinct mechanism of action that has yet to be fully characterized.

Some T2Rs, including T2R5, are upregulated in malignant tissues compared to normal tissue (13), but their functional roles in cancer remain underexplored (13). It is unknown, for example, if all T2Rs in HNSCC cells can activate apoptosis. Given that T2R14 and T2R4 activation causes cell death in HNSCC and phendione is a relatively specific and potent T2R5-agonist, we hypothesized that phendione would induce cell death and decrease cell viability in HNSCC via T2R5. We investigated the role of T2R5 in HNSCC to evaluate the potential of phendione or other T2R5 agonists as chemotherapeutic agents. Using *in vitro* and *ex vivo* models, we demonstrate that phendione induces intracellular Ca^2+^ responses, reduces cell viability, and promotes apoptosis through T2R5-dependent mechanisms. Furthermore, we explore the association between T2R5 expression and patient survival, offering new insights into the therapeutic potential of T2Rs in HNSCC.

## Materials and Methods

### Cell Culture

Human cell lines SCC47 (lateral tongue, HPV+) (UM-SCC-47; Millipore SCC071, RRID:CVCL_7759), FaDu (hypopharynx, HPV-) (ATCC HTB-43, RRID:CVCL_1218), RPMI 2650 (nasal septum, HPV-) (ATCC CCL-30, RRID:CVCL_1664), SCC4 (tongue, HPV-) (ATCC CRL-1624, RRID:CVCL_1684), and SCC90 (base of tongue, HPV+) (ATCC CRL-3239, RRID:CVCL_1899) were cultured submerged in high glucose DMEM (Corning; Glendale, AZ, USA) supplemented with 10% FBS (Genesee Scientific; El Cajon, CA, USA), 1% penicillin/streptomycin mix (P/S) (Gibco; Gaithersburg, MD, USA), and 1% nonessential amino acids (Gibco). Primary gingival keratinocytes (PCS-200-014) were grown submerged with Dermal Cell Basal Medium (PCS-200-030) with Keratinocyte Growth Kit (PCS-200-040) from ATCC.

Tissue was acquired from HNSCC patients undergoing oncologic surgeries after written informed consent and approval of the University of Pennsylvania Institutional Review Board (IRB #417200). Patient ages ranged from 51.1 to 77.0 years; all included participants were male. Patients were only included if they were undergoing curative-intent surgery for primary HNSCC and had adequate additional tissue available for research. Patients were excluded if they had pathologies other than HNSCC. No patient or sample attrition occurred during the study. Sex of patients was recorded but was not used as a variable in analysis as all patients were male. All specimens processed were included in the final analysis. *Ex vivo* human tumor samples were sectioned into 300 µm thick tumor slice cultures using a vibratome (Precisionary Compresstome® Vibratome VF-510-0Z) with the following settings: oscillations 6, speed 3.

### Immunofluorescence

Cells were grown on chambered glass coverslips (CellVis) and fixed with 4% paraformaldehyde in PBS and subsequently blocked and permeabilized in buffer containing 1% BSA, 0.1% Triton X-100 in PBS. Primary T2R5 antibodies (Bioss) were applied at 1:250 dilution and incubated overnight at 4°C. Secondary antibodies, Alexa Fluor-546 (Thermo Scientific), were incubated for 2 hours at a 1:500 dilution. DAPI was used as a nuclear counterstain using Fluoromount Mounting medium (Thermo Fisher Scientific). Images were captured using the Olympus Live Cell imaging System with 40x objectives. For *ex vivo* tumor slices, phalloidin conjugated to Alexa fluor-488 (Life Technologies) was used as an actin counterstain. Images were processed through Fiji version of ImageJ software (RRID:SCR_002285) (38).

### Reverse Transcription Quantitative PCR (RT-qPCR)

Cells were lysed in TRIzol (Thermo Fisher Scientific), and RNA was isolated and purified using the Direct-zol RNA kit (Zymo Research). Reverse transcription was performed with High-Capacity cDNA Reverse Transcription Kit (Thermo Fisher Scientific), and *TAS2R5* expression was quantified using TaqMan qPCR assays.

Ubiquitin C (UBC) was selected as the housekeeping gene for normalization, due to its stable expression in cancer cells (39).

### siRNA Transient Knockdown TAS2R5

SCC47 cells were transfected with *siTAS2R5* (hs.Ri.TAS2R5.13) or *siScramble* control (IDT DsiRNA TriFecta kit) with RNAiMAX lipofectamine reagent (Thermo Fisher Scientific) and Opti-MEM (Gibco). Suspended transfections were performed on day 1, with additional transfection on day 3. Knockdown efficiency was validated by western blot, showing sufficient T2R5 protein knockdown compared to control.

### Live Cell Ca^2+^ Imaging

Cells were loaded with 5 µM of Fluo-4-AM (Thermo Fisher Scientific) for 1 hour at room temperature in the dark. Imaging was performed in Hank’s Balanced Salt Solution (HBSS) buffered with 20 mM HEPES (pH 7.4) containing 1.8 mM Ca^2+^. Cells were imaged on 8-well glass bottom chamber slides using Olympus IX-83 microscope (20x 0.75 NA PlanApo objective) with FITC filters (Sutter Lambda LS), Orca Flash 4.0 sCMOS camera (Hamamatsu, Tokyo, Japan), Metamorph (Molecular Devices), and XCite 120 LED Boost (Excelitas Technologies). Ligands (phendione, 1,10-phenanthroline, procyanidin trimer C2) were dissolved in HBSS and added in real-time during imaging. Final DMSO concentration did not exceed 0.1%. Images were analyzed through Fiji ImageJ software (38).

For zero-extracellular Ca^2+^ (“0-Ca^2+^”) conditions, cells were loaded with Fluo-4-AM in HBSS, agonists tested were dissolved in Ca^2+^-free HBSS with 10 mM EGTA (pH = 7.4). At the time of ligand injection, cells were bathed in 60 µL of Ca^2+^ containing HBSS and agonist was added in a volume of 300 µL and a final concentration of ∼8.3 mM EGTA. We previously calculated that this is equal to final free Ca^2+^ concentration of 2 nM using MaxChelator (Chris Patton, Stanford University, RRID:SCR_000459) (14).

For ER Ca^2+^ imaging, pCMV-ER-LAR-GECO1 (Addgene_61244) (40) was transfected into cells using Lipofectamine 3000 (Thermo Fisher Scientific) 24 hours prior to imaging. Gα_q_ inhibitors YM254890 (FUJIFILM Wako Chemicals) and FR900359 (Sapphire North America) were incubated at 1 µM and 10 µM, respectively, for 1 hour at room temperature in the dark. Pertussis toxin (Tocris) (50 ng/mL) was incubated overnight.

### Cell Viability: Crystal Violet Assay

Cells were treated with phendione in high-glucose DMEM +10% FBS, +1% P/S, +1% NEAA for 24 hours at 37° C in a humidifier incubator at 5% CO_2_. Remaining adherent cells were stained with 0.1% crystal violet, washed, dried, and dissolved in 30% acetic acid. Absorbance at 590 nm was measured using a Tecan plate reader (Tecan Spark 10M; Mannedorf, Switzerland) and normalized to untreated controls.

For *ex vivo* tumor slices, viability changes at 24 hours were assessed using CellTiter 96 Aqueous One solution MTS Cell Viability dye (Promega). Adjacent tumor slices were incubated in phenol-free, high glucose DMEM +10% FBS, +1%P/S, +1% NEAA with or without experimental treatments, MTS cell viability dye was added at 22 hours, to both tumor slices and blanks, and incubated for an additional 2 hours for total duration of 24 hours. Adjacent tumor slices have previously been shown to be metabolically similar, serving as technical replicates (41). After incubation period, 50 µL of media from tumor slices and blanks were transferred to a new well and absorbance at 490 nm was measured using Tecan plate reader. Results were normalized to untreated controls for each patient.

### Apoptosis Assays

Cells were seeded into 24-well black glass-bottom plates (CellVis) to achieve ∼70% confluency at the time of the experiment. Treatments were prepared in phenol-free, high glucose DMEM +10% FBS, + 1% P/S, and + 1% NEAA. CellEvent Caspase 3/7 indicator dye (Thermo Fisher Scientific) was added per manufacturer’s instructions. Images were acquired every 25 minutes for 24 hours using a Tecan Plate reader with 5% CO_2_ at 37° C. Representative images were taken on microscope described above.

### The Cancer Genome Atlas (TCGA) Analyses

*TAS2R5* expression data were obtained from the TCGA, TARGET, GTEx combined cohort via UCSC Xena (https://xenabrowser.net/). DESeq2-standardized counts (RRID:SCR_015687) were used to compare *TAS2R5* expression levels across datasets. Kaplan-Meier survival analyses were performed using Xena to evaluate the correlation of *TAS2R5* expression with overall and disease-specific survival. For comparison of *TAS2R5* expression between tumor and normal samples, log2-transformed DESeq2 standardized counts were utilized. Graphs were generated in GraphPad Prism (RRID:SCR_002798, San Diego, CA, USA).

### Statistical Analyses

Statistical analyses were conducted using GraphPad Prism (San Diego, CA, USA). T-tests were used for two-group comparisons, while one-way ANOVA was applied for comparisons across multiple groups. Sample sizes were chosen based on prior publications demonstrating sufficient power to detect treatment effects in similar models; no formal power analysis was performed. Statistical tests used are indicated in the figure legends, with significance indicated as follows: P < 0.05 (*), P < 0.01 (**), P < 0.001 (***), (****) P < 0.0001. Error bars represent mean ± SEM unless otherwise indicated.

### Data Availability

All data generated in this study are shown in the manuscript. The raw data points used to generate graphs and figures are available upon request from the corresponding author.

## Results

### Endogenous Expression of T2R5 in HNSCC Cell Lines and Patient Derived Tumor Slices

The bitter taste receptor T2R5 has been implicated in various physiological and pathological processes (29,42,43), but its expression in HNSCC remains underexplored. To determine whether *TAS2R5* is endogenously expressed in HNSCC, we assessed its expression across multiple cell lines representing both HPV+ and HPV− HNSCC subtypes. We found that *TAS2R5* is variably expressed in both HPV+ (SCC47) and HPV-(RPMI 2650, SCC4, FaDu) HNSCC cell lines, as detected by RT-qPCR (**Figure 1A**). Comparisons to non-malignant primary gingival keratinocytes showed lower expression of *TAS2R5* compared with SCC47, RPMI 2650, and SCC4 cells but no significant difference compared with FaDu cells.

**Figure 1:**
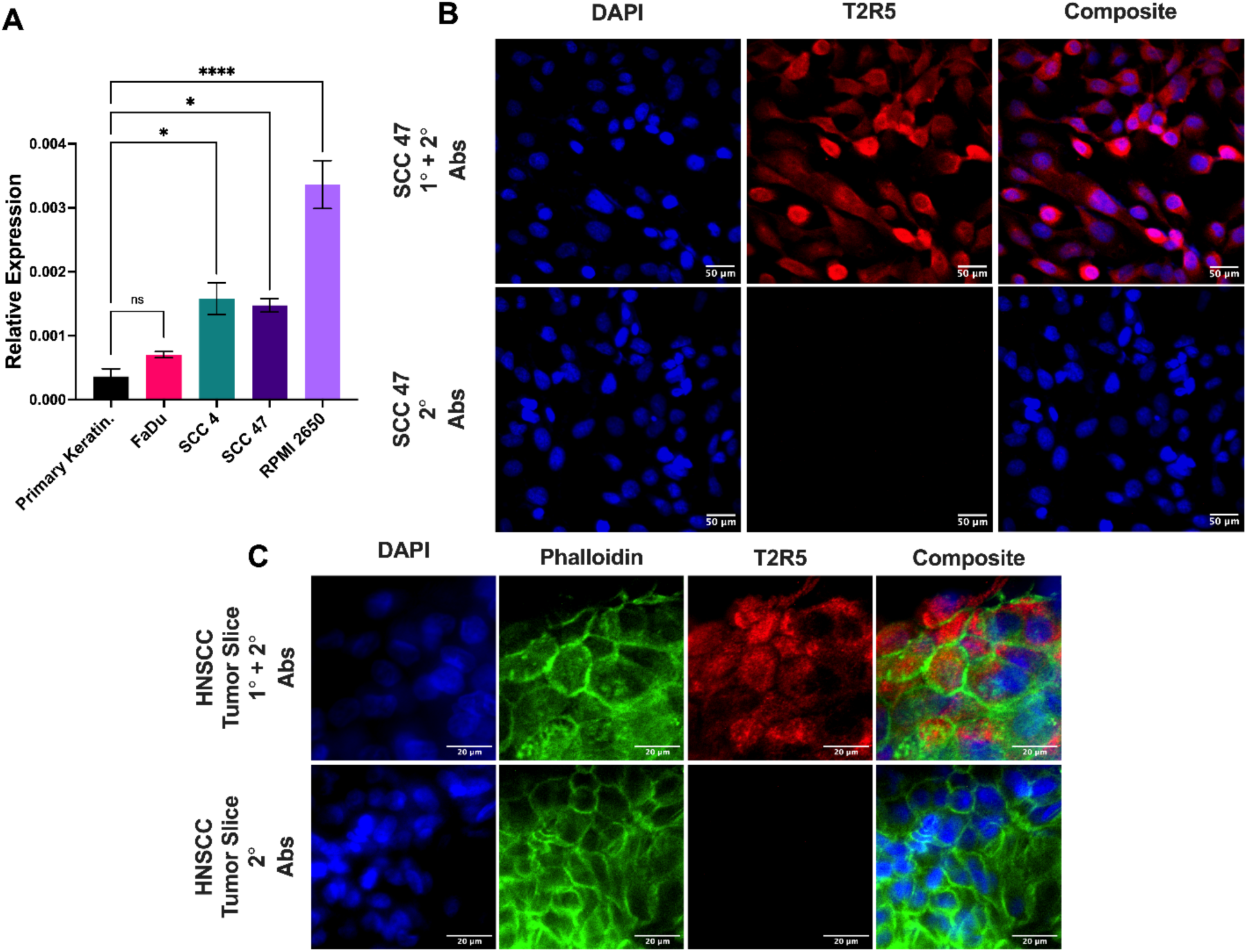
***TAS2R5* is expressed in HNSCC cell lines and patient-derived tumors. A** Relative mRNA expression of *TAS2R5* was measured in primary keratinocytes and compared to HNSCC cell lines FaDu, SCC4, SCC47, and RPMI2650 by qPCR normalized to ubiquitin C *(UBC)* housekeeping gene, (mean ± SEM, n > 3 separate passages). **B** T2R5 protein expression measured via immunofluorescence in SCC47 (DAPI = nucleus). **C** *Ex vivo* patient-derived tumor slice from oral tongue HPV-HNSCC showing diffuse T2R5 staining (DAPI = nucleus; phalloidin = (green, actin). Each antibody was compared to secondary only control at the same microscope settings. Significance by 1-way ANOVA with Dunnett’s multiple comparison posttest. *p < 0.05, **p < 0.01, ***p < 0.001; ns, no statistical significance.

Immunofluorescence using a T2R5 antibody revealed diffuse staining in both SCC47 (**Figure 1B**) and RPMI 2650 (**Supp. Figure 1**) cells. These findings suggest that T2R5 is endogenously expressed in HNSCC cell lines. Immunofluorescence of patient derived tumor samples (**Figure 1C**) also showed T2R5 staining, suggesting that T2R5 is present within HNSCC tumors.

### Stimulation with Phendione Induces a Sustained Intracellular Ca^2+^ Response in HNSCC

Previous studies have identified 1,10-phenanthroline as the only compound among ∼100 bitter ligands capable of activating T2R5 in heterologously expressed HEK-293T cells (26). In human airway smooth muscle cells, its derivative, 1,10-phenanthroline-5,6-dione (phendione), induces Ca^2+^ responses at much lower concentrations, suggesting that phendione is a more potent T2R5 agonist (29). To test if phendione stimulation elicits a Ca^2+^ response in HNSCC, live cell imaging was performed on cells loaded with the Ca^2+^ indicator dye Fluo-4.

In SCC47 cells, phendione stimulation at concentrations as low as 1.4 µM induced a significant intracellular Ca^2+^ response that was sustained for ≥30 minutes (**Figure 2A-B**). Similar responses were observed in RPMI 2650, FaDu, and SCC4 cell lines with 1.4 and 142 µM phendione (**Figure 2C-E**), while SCC90 cells demonstrated a significant response only at 142 µM (**Figure 2F**). In primary gingival keratinocytes, phendione also elicited significant Ca^2+^ responses (**Figure 2G-H**), but the response was oscillatory rather than sustained over time (**Figure 2A**). Notably, in SCC47, the sustained Ca^2+^ responses persisted for ≥5 hours and resulted in morphological changes. (**Figure 2I-J)**. These findings highlight differences in T2R5-activated Ca^2+^ response kinetics between malignant and non-malignant cells.

**Figure 2:**
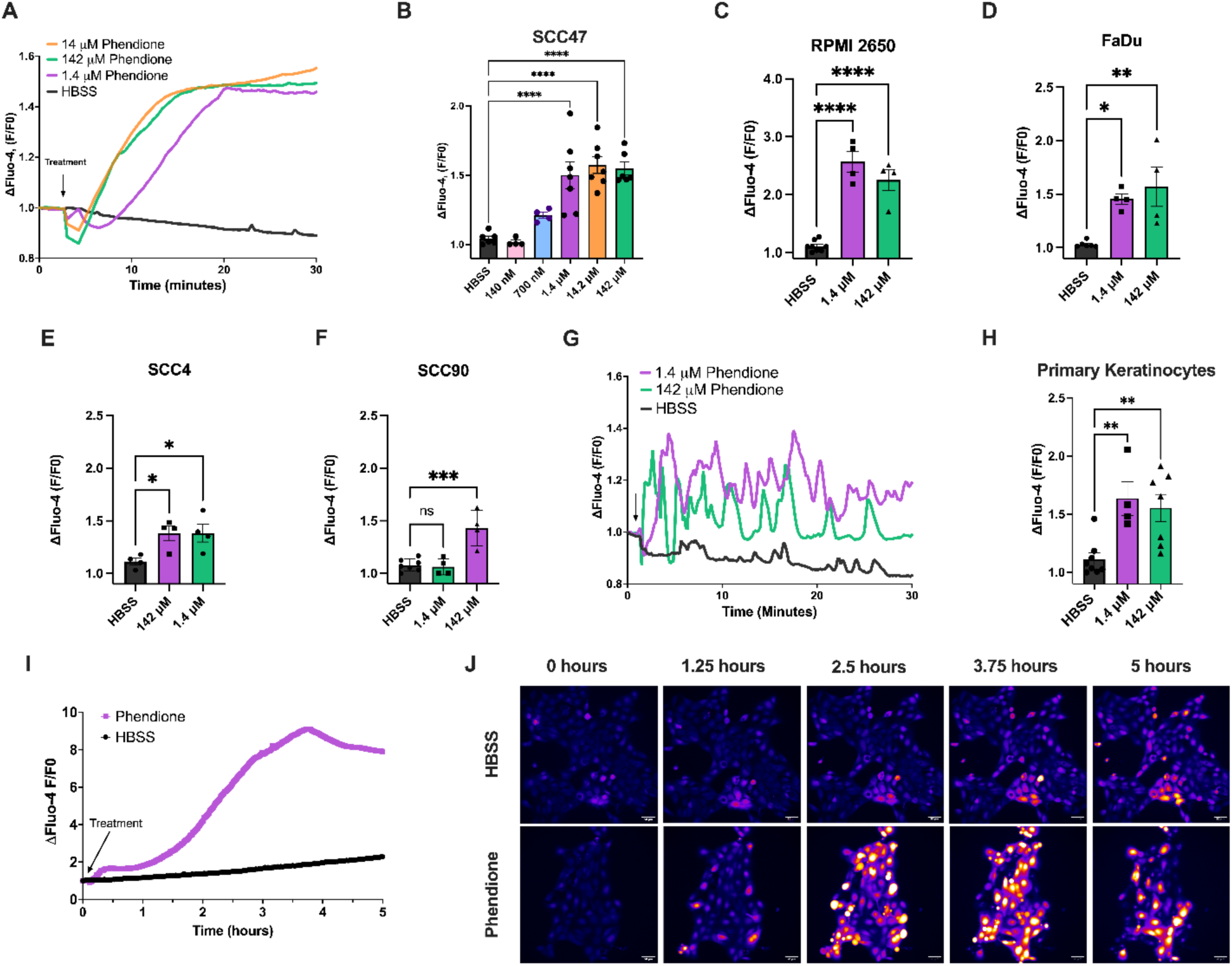
Phendione stimulation results in sustained, intracellular Ca^2+^ responses. HNSCC cell lines were loaded with Ca^2+^ binding dye Fluo-4-AM and stimulated with T2R agonist phendione. **A-B** SCC47 Ca^2+^ responses over time (A, minutes) and peak (B, mean ± SEM) after phendione stimulation. **C-F**: Peak Ca^2+^ response after phendione stimulation for RPMI2650 (C), FaDu (D), SCC4 (E), and SCC90 (F). **G-H:** Primary keratinocyte Ca^2+^ responses over time (C, minutes) and peak (D, mean ± SEM) after phendione stimulation. Significance by 1-way ANOVA with Dunnett’s multiple comparison posttest comparing HBSS to each agonist concentration. **I-J:** Extended evaluation of SCC47 Ca^2+^ responses to phendione demonstrate a steady increase that peaks around 4 hours. *p < 0.05, **p < 0.01, ***p < 0.001, ****p<0.0001; ns, no statistical significance.

To assess the source of Ca^2+^ mobilization, phendione stimulation was repeated under zero extracellular Ca^2+^ (0-Ca^2+^) conditions, achieved by chelating extracellular Ca^2+^ with EGTA. Under these conditions, phendione induced an immediate but transient Ca^2+^ peak that was not sustained, unlike the response observed under normal extracellular Ca^2+^ (1.8 mM Ca²⁺; **Figure 3A-B**). This indicates that the initial Ca^2+^ response primarily arises from intracellular Ca^2+^ stores, likely the ER.

**Figure 3:**
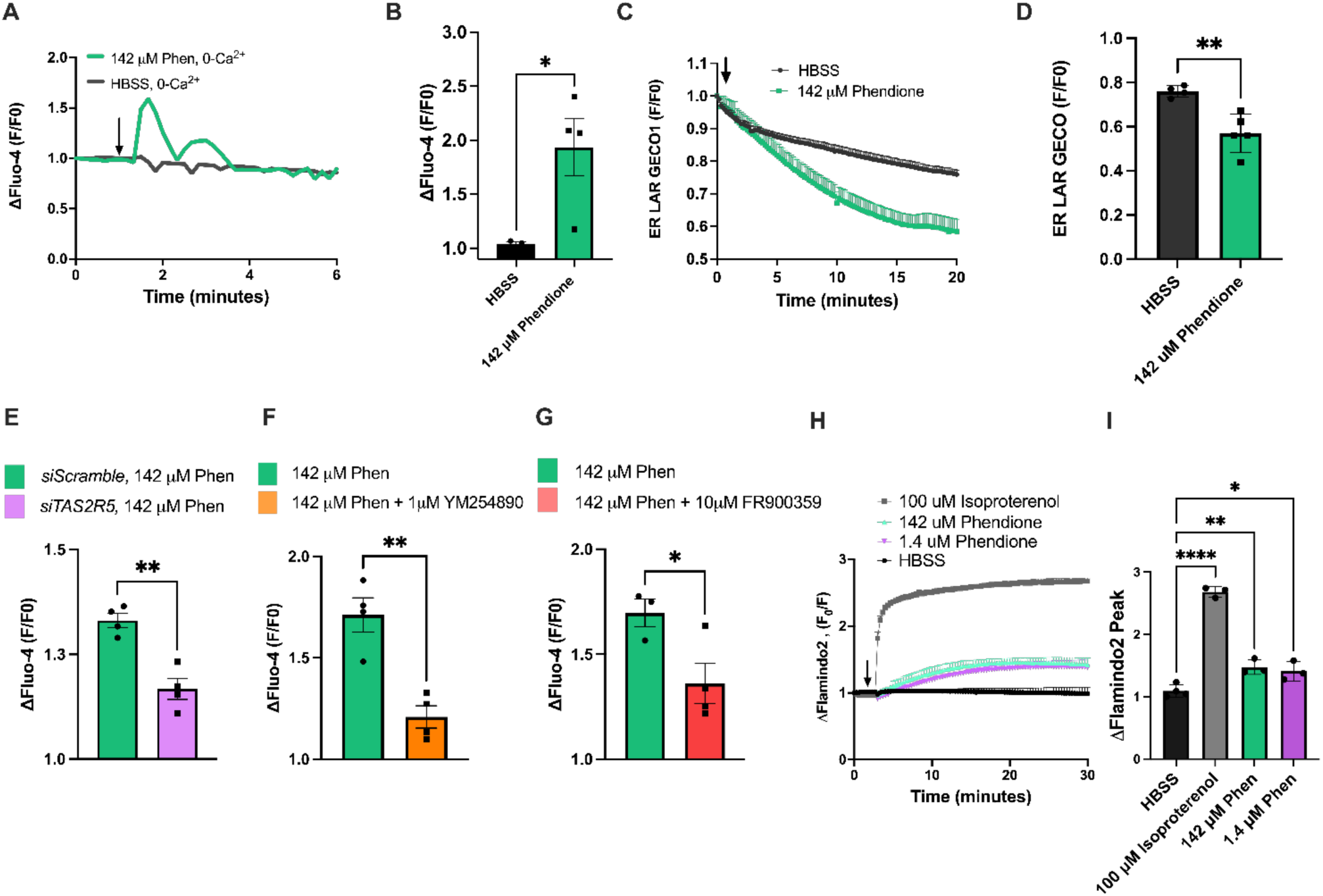
Phendione intracellular Ca^2+^ responses are T2R5 dependent. **A-B** To assess the source of Ca^2+^ mobilization, phendione stimulation was repeated under 0-extracellular Ca^2+^ conditions (0-Ca^2+^), achieved by chelating extracellular Ca^2+^ with EGTA. Phendione induced an immediate, but transient Ca^2+^ peak that was not sustained, suggesting that the initial Ca^2+^ response primarily arises from intracellular Ca^2+^ stores. **C-D** The role of ER Ca^2+^ stores was evaluated by transfecting SCC47 cells with the fluorescent ER Ca^2+^ biosensor pCMV ER LAR GECO1. Phendione stimulation caused a steady and significant depletion of ER Ca^2+^compared to HBSS control (C). Endpoint Ca^2+^ levels in the ER after 20 minutes were significantly lower in the phendione condition versus HBSS (D). **E** SCC47 transient knockdown of *TAS2R5* was generated using siRNA (*siTAS2R5*) and compared to nontargeting human siRNA (*Scramble*). Peak Ca^2+^ responses (mean ± SEM) were significantly reduced for *siTAS2R5* compared to *siScramble* suggesting T2R5-dependent signaling. **F-G** SCC47 cells were loaded with Ca^2+^ binding dye, Fluo-4, and treated with Gαq inhibitors YM254890 (F) and FR900359 (G). Gαq inhibitors significantly blunted the peak Ca^2+^ responses to phendione compared to untreated controls (mean ± SEM), suggesting Gα involvement. **H-I** Cyclic AMP (cAMP) production was assessed using fluorescent biosensor pcDNA3.1-pFlamindo2. In contrast to most bitter agonist signaling to Gα_i/o_ which leads to decreased cAMP, stimulation of SCC47 cells with phendione resulted in an increase in cAMP levels compared to the HBSS control. Isoproterenol (positive control) induced a robust increase in cAMP production. Significance by unpaired t-test. Significance by 1-way ANOVA with Dunnett’s multiple comparison posttest. *p < 0.05, **p < 0.01, ***p < 0.001; ns, no statistical significance.

The role of ER Ca^2+^ stores was further evaluated by transfecting SCC47 cells with the fluorescent ER Ca^2+^ biosensor ER-LAR-GECO1 (40). Phendione stimulation caused a steady and significant decrease of ER Ca^2+^ compared to HBSS control, as shown by a gradual decrease in fluorescence (**Figure 3C**). Endpoint Ca^2+^ levels in the ER after 20 minutes were significantly lower in the phendione condition versus HBSS control (**Figure 3D**). Together, these data suggest that phendione mobilizes Ca^2+^ from ER stores, a typical characteristic of T2R activation (7,44,45). However, the sustained Ca^2+^ response observed in SCC47 cells was absent under 0-Ca^2+^ conditions, indicating a requirement for extracellular Ca^2+^ influx during prolonged stimulation.

To evaluate Ca^2+^ responses to other previously reported T2R5-selective agonists, SCC47 cells were stimulated with 1,10-phenanthroline and procyanidin trimer C2 (procyanidin) (26,46–48). Significant dose-dependent Ca^2+^ responses were observed for both procyanidin and 1,10-phenanthroline (**Supp. Figures 2**). These data provide further evidence that T2R5 agonism can lead to significant, sustained Ca^2+^ responses in HNSCC cell lines.

### Phendione Acts on Endogenously Expressed T2R5 Likely Coupled to Gα_q_ in HNSCC

To determine whether the phendione-induced Ca^2+^ response is mediated by T2R5, transient knockdown of *TAS2R5* was performed using siRNA (*siTAS2R5*). This resulted in a significant reduction in Ca^2+^ response compared to cells transfected with a nontargeting siRNA control (*siScramble*), indicating that the Ca^2+^ response is primarily T2R5-dependent (**Figure 3E**). Successful T2R5 protein knockdown was confirmed with Western blot (**Supp. Figure 3**). Furthermore, these findings align with published data demonstrating the selectivity of 1,10-phenanthroline (parent compound to phendione) for T2R5 over other T2Rs (26).

To explore the role of G-protein coupling in the phendione-induced Ca^2+^ response, live cell imaging with Fluo-4 was conducted in the presence of specific G-protein inhibitors. Two Gα_q_ inhibitors, YM254890 and FR900359 (49,50), significantly blunted the Ca^2+^ response to phendione compared to untreated controls (**Figure 3F-G**). In contrast, treatment with pertussis toxin (PTX), a Gα_i/o_ inhibitor that we showed reduces T2R14-mediated responses (14), did not significantly decrease the Ca^2+^ response (**Supp. Figure 4A**). Additionally, the T2R14 antagonist LF1 (51) did not attenuate the Ca^2+^ response to phendione (**Supp. Figure 4B**), further supporting that phendione targets T2R5 and not other bitter taste receptors such as T2R14.

To investigate secondary messenger signaling, cyclic AMP (cAMP) production was assessed using fluorescent biosensor pFlamindo2 (52), which enables real-time visualization of cAMP dynamics. T2Rs coupled to Gα_i/o_ can inhibit adenylate cyclase, leading to a decreased in cAMP (53,54). However, stimulation of SCC47 cells with phendione resulted in a steady and significant *increase* in cAMP levels compared to the HBSS control (**Figure 3H-I**). Isoproterenol, a β-adrenergic agonist, was used as a positive control to induce cAMP production (55,56). These results support that phendione does not activate Gα_i/o_ but rather signals through a distinct GPCR pathway, likely involving Gα_q_. Ca^2+^ responses with stimulation by another T2R5 agonist, procyanidin, was also reduced by Gα_q_ inhibitor YM254890, further suggesting that T2R5 couples to Gα_q_ (**Supp Figure 2C**).

### Phendione Exposure Reduces Cell Viability in a Dose-Dependent Manner

Phendione was previously identified as a potential chemotherapeutic with cytotoxic effects (57). To evaluate its impact on HNSCC cells via T2R5, crystal violet assays were performed to measure cell viability following 24-hour phendione exposure. A dose-dependent reduction in cell viability was observed across HNSCC cell lines (**Figure 4**). Concentrations as low as 140 nM caused a significant decrease in cell viability in SCC47 and RPMI 2650 (**Figure 4A-B**), while SCC4 and FaDu viability was significantly reduced at 1.42 µM (**Figure 4C-D**).

**Figure 4:**
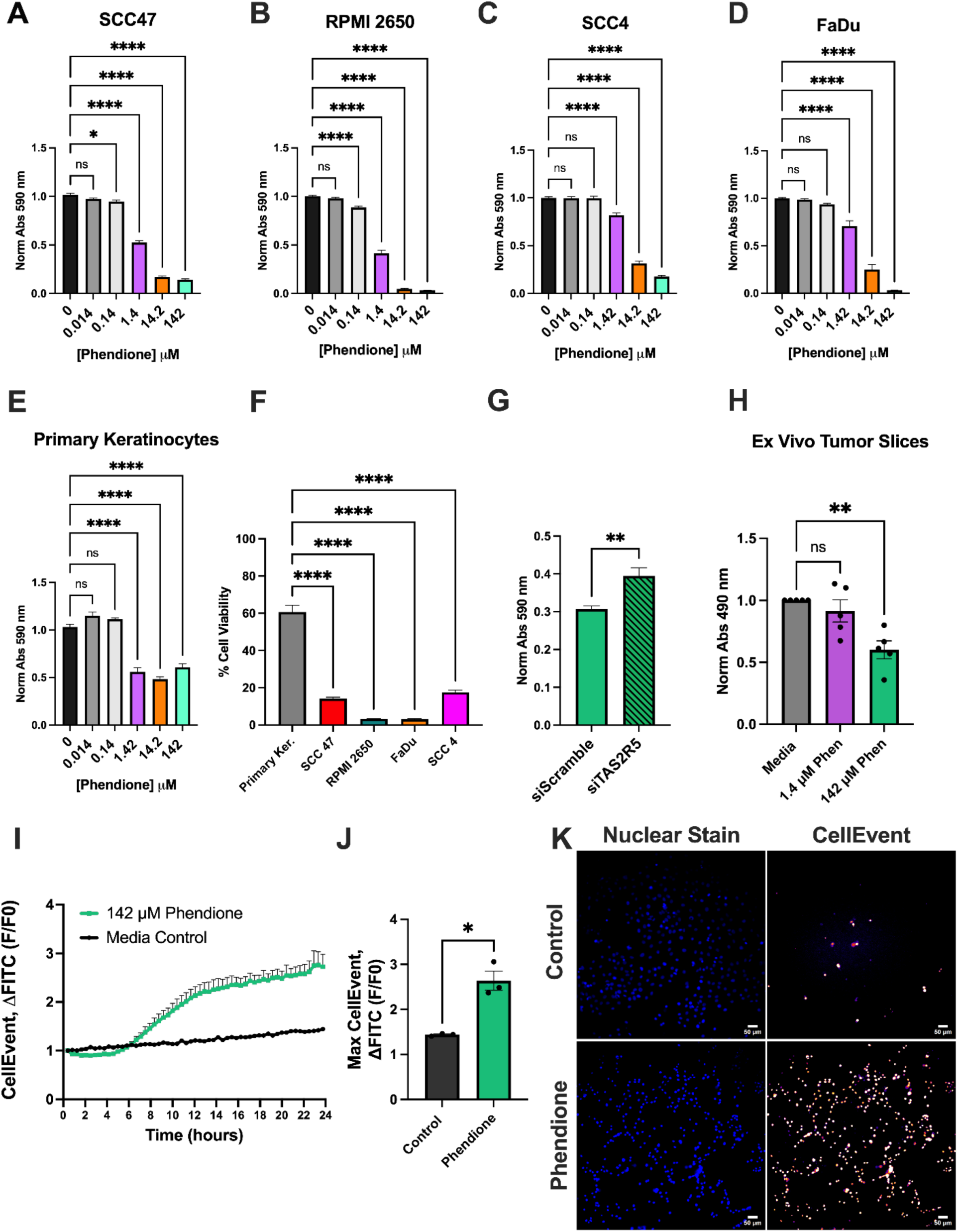
T2R5 stimulation leads to HNSCC apoptosis and reduced cell viability. **A-H** Cell viability was measured following 24-hour phendione exposure using crystal violet and measured at 590 nm absorbance. Dose dependent reductions in cell viability were observed in cancer cell lines (A-D) and primary keratinocyte (E; mean ± SEM, one-way ANOVA Dunnett’s multiple comparison test). **F** The percent viability after 142 µM phendione exposure was significantly lower for cancer cell lines compared to primary keratinocytes (mean ± SEM, one-way ANOVA Dunnett’s multiple comparison test). **G** SCC47 *siTAS2R5* transient knockdown exhibited less robust decreases in cell viability after phendione treatment compared to control *siScramble* (mean ± SEM, unpaired t-test with Welch’s correction). **H** Viability of HNSCC tumor slices from n = 5 patients indicates significant reduction after 24 hours of treatment with 142 µM phendione. No significant reduction is observed at 1.4 µM phendione. Significance by 1-way ANOVA with Dunnett’s multiple comparison posttest. **I-K** To evaluate if phendione induces apoptosis, SCC47 cells were stained with CellEvent, a dye that fluoresces upon activation of caspase 3/7 cleavage. Phendione increased fluorescence compared to controls over 24 hours (I, J; mean ± SEM; unpaired t-test). **K** Representative images at 24 hours with Hoechst dye (counter nuclear stain) and CellEvent dye in control and phendione condition. *p < 0.05, **p < 0.01, ***p < 0.001; ns, no statistical significance.

In primary gingival keratinocytes, phendione also caused a dose-dependent reduction in cell viability (**Figure 4E**). However, keratinocytes showed greater resistance to phendione at 142 µM phendione compared to HNSCC cell lines (**Figure 4F**), suggesting that, although phendione is not entirely cytoselective for cancer cells, normal cells may exhibit greater resistance to its cytotoxic effects. Calculated IC_50_ values for phendione in HNSCC cell lines range from ∼0.8 – 4.8 µM (**Supp. Figure 5**), consistent with previously reported EC_50_ values in other cell lines (29,58).

To investigate whether the reduction in cell viability is mediated through T2R5, SCC47 cells with transient T2R5 knockdown (*siTAS2R5*) were evaluated. Cells transfected with *siTAS2R5* demonstrated improved viability at 142 µM phendione compared to cells transfected with non-targeting *siScramble* controls (**Figure 4G**). This indicates that phendione acts via T2R5 to decrease HNSCC cell viability.

To further assess phendione’s effects, *ex vivo* tumor slice cultures derived from HNSCC patients were exposed to 1.4 and 142 µM phendione for 24 hours. Cell viability was measured using MTS dye, with results normalized to controls. Phendione significantly reduced cell viability at 142 µM (**Figure 4H**). The demographic information for the HNSCC tumor samples can be found in **Supp. Table 1**. This *ex vivo* data corroborates findings from HNSCC cell lines, further supporting anti-tumor cytotoxic effects of phendione and potential translational applications to patients.

To evaluate if phendione induces apoptosis, SCC47 cells were stained with CellEvent, a dye that fluoresces upon activation of caspase 3/7 cleavage. Phendione (142 µM) significantly increased fluorescence compared to controls, indicative of caspase 3/7 activation (**Figure 4I-K**). Fluorescence began increasing at ∼6 hours post-treatment and steadily rose over the 24-hour period (**Figure 4I**). These results suggest that phendione induces apoptosis in HNSCC cells through caspase 3/7 activation, consistent with prior reports of similar mechanisms involving other T2Rs in HNSCC (13,14).

### TAS2R5 Expression in HNSCC Tumors is Associated with Long-Term Patient Survival

To evaluate the clinical implications of *TAS2R5* expression in HNSCC, TCGA data were analyzed to compare patients with low versus high *TAS2R5* expression levels based on bulk RNA-sequencing of tumor samples. Kaplan-Meier survival analyses demonstrated improved 5-year overall survival and disease-specific survival for cases with high *TAS2R5* expression compared to low *TAS2R5* expression (**Figure 5A-B**) which was also observed at 10 years (**Supp. Figure 6**). These findings suggest a potential endogenous tumor-suppressive role of *TAS2R5* in regulating tumor proliferation in HNSCC patients.

**Figure 5:**
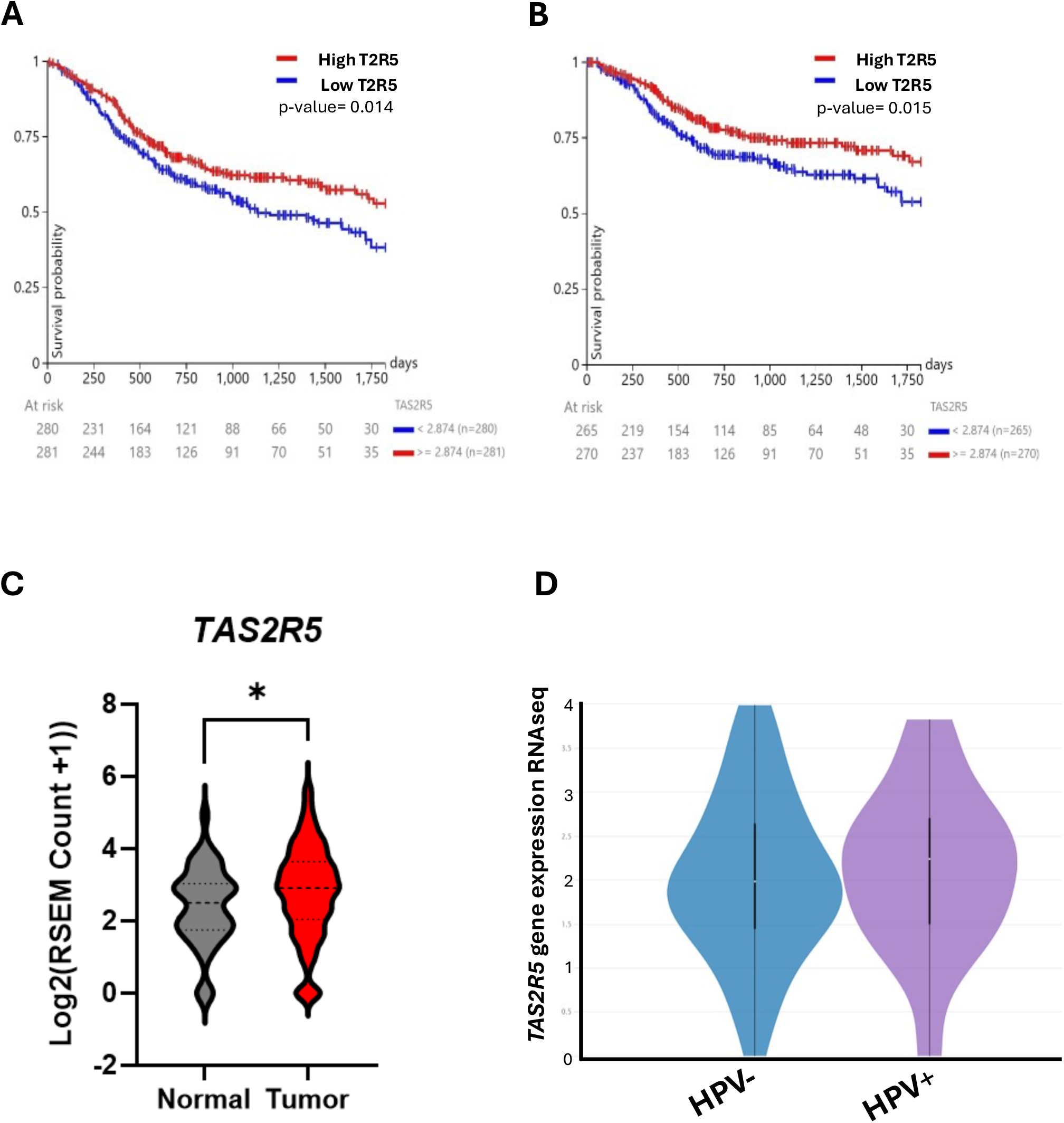
Increased *TAS2R5* expression is associated with improved survival. A-B. Using bulk RNA-sequencing data from The Cancer Genome Atlas (TCGA), HNSCC patients were divided into groups based on high and low *TAS2R5* expression. Kaplan– Meier survival analysis demonstrated improved 5-year overall survival (A) and disease-specific survival (B) for cases with high *TAS2R5* expression compared to low *TAS2R5* expression (p = 0.014 and 0.015 by log-rank test, respectively). **C** Comparison of log2 transformed expression levels indicated a slight increase in *TAS2R5* expression in tumor compared to adjacent normal tissue in HNSCC patients. *p < 0.05. **D** Comparison of log2 transformed expression levels of *TAS2R5* indicate no significant difference between HPV+ and HPV– HNSCC tumors (n= 113, Welch’s t-test p = 0.61).

Our prior study highlighted differential expression of *TAS2Rs* between tumor and normal tissues, with *TAS2R5* showing higher expression in tumor samples (13). To evaluate this further, *TAS2R5* expression levels in TCGA were compared between tumor and adjacent normal tissues in HNSCC patients. Results suggest a slight but significant increase in *TAS2R5* expression in tumor tissue (**Figure 5C**). Previously, we demonstrated that *TAS2R14* expression was increased in HPV+ HNSCC tumors (14). To investigate if *TAS2R5* expression is similarly influenced by HPV status, TCGA data was stratified into HPV+ and HPV− HNSCC tumors. No significant differences in *TAS2R5* expression were observed between the two groups (**Figure 5D**). Unlike *TAS2R14*, *TAS2R5* expression is not specifically upregulated in HPV+ tumors.

## Discussion

Bitter taste receptors (T2Rs), a subclass of GPCRs, have increasingly been recognized for their crucial extra-oral roles in both normal (10,27) and malignant tissues (12,59). Recent studies have highlighted their therapeutic potential in HNSCC (13–15,60), with ongoing research that includes a phase I clinical trial led by our group to evaluate T2R14-agonist lidocaine as a neoadjuvant treatment for HNSCC (22). While T2R14 has been explored for its cytotoxic effects in HNSCC (13,14), the role of T2R5 – a highly selective receptor for specific bitter ligands (27) – remained unexplored until now. T2R5 has been implicated in extra-oral functions across multiple tissues, including the brain, uterus, skin, and lung (29,43,61,62), but its role in cancer had not been defined.

We demonstrate that phendione, a high-affinity T2R5 agonist, exerts cytotoxic effects in HNSCC via T2R5-dependent mechanisms. Phendione, a derivative of 1,10-phenanthroline (29), was previously noted to exhibit anticancer activity in epithelial cells (57). Unlike traditional metal chelators like cisplatin, phendione induces anti-cancer effects without causing DNA breaks (35), suggesting alternative cytotoxic pathways. Our data suggest that phendione activates T2R5, leading to sustained Ca^2+^ signaling, reduced cell viability, and apoptosis. These data may indicate T2R5 as a potential viable target for HNSCC therapies.

### T2R5 Expression in HNSCC

We identified endogenous expression of T2R5 in HNSCC, with varying levels across different cell lines. Primary keratinocytes exhibited the lowest T2R5 expression, potentially explaining their relative resistance to phendione-induced cytotoxicity compared to malignant cells. *Ex vivo* tumor slices also demonstrated T2R5 expression, supporting the clinical relevance of T2R5 as a therapeutic target. The observed differential expression between malignant and normal tissues suggests that T2R5 activation could selectively target cancer cells while minimizing off-target effects in normal tissues.

### T2R5 induces Ca^2+^ Mobilization in HNSCC cells

Phendione stimulation induced a sustained intracellular Ca^2+^ response in HNSCC cells. This contrasts with the transient responses observed in primary keratinocytes and other T2R subtypes in HNSCC cells, like T2R14, where Ca^2+^ peaks more transiently (14,53). Sustained Ca^2+^ signaling has been implicated in mitochondrial dysfunction, ROS production, and caspase activation, all of which contribute to apoptosis (63–66). This could be particularly relevant for tumors with high T2R5 expression that are stimulated with a T2R5-agonist to induce or sensitize malignant cells to undergo apoptosis. Additionally, the responses to other T2R5 agonists, like 1,10-phenanthroline and procyanidin trimer C-2, further supports the role of T2R5 in the Ca^2+^ mobilization observed with phendione. Procyanidin trimer C-2, a potent polyphenol in fruits and berries (46), has been reported to exert preventative and anti-cancer effects in HNSCC (67–69). While some reports have evaluated the role of dietary intake of plant polyphenols in HNSCC (70), future stuides could evaluate the effects of polyphenols that specifically stimulate T2Rs.

Interestingly, our data suggest that phendione/T2R5 activates Ca^2+^ signaling through Gα_q_-mediated pathways. We have showed that denatonium, a highly bitter agonist that activates many T2Rs, is also inhibited by Gα_q_-inhibitors (13). While some T2Rs are blocked by Gα_i/o_ inhibitors like pertussis toxin (14), T2R5-agonists are not. This G-protein coupling difference emphasizes the complexity of T2R signaling and opens new avenues for targeted therapies that can leverage specific signaling pathways. GPCRs as a whole make up the largest portion of currently approved therapeutics (71), and many current FDA-approved therapeutics are predicted to activate bitter taste receptors (23,51,72). It is thus important to characterize tissue specific T2R signaling for drug development, repurposing, and off-target effects.

### Cytotoxic Effects of T2R5 activation by Phendione

The dose-dependent reduction in cell viability observed with phendione treatment highlights its therapeutic potential in HNSCC. While primary keratinocytes also exhibited reduced viability, their relative resistance compared to malignant cells points to a role for differential T2R5 expression in mediating cytotoxic effects. This suggests that phendione or similar T2R5 ligands could be leveraged as selective chemotherapeutic agents, particularly in tumors with elevated T2R5 expression.

The induction of apoptosis further reinforces the potential specificity of T2R5-mediated cytotoxicity. Apoptosis is a critical mechanism for eliminating cancer cells while minimizing damage to surrounding tissues (73–75) and targeting T2R5 may provide an opportunity to exploit this pathway in HNSCC. Importantly, this approach may complement existing therapies, as apoptosis induction is often impaired in resistant tumors (76,77). Future work is needed to explore the potential of phendione or other T2R5 agonists to synergize with chemotherapeutic agents such as cisplatin, which relies on DNA damage pathways, or BAY-876, which targets GLUT1 and has demonstrated efficacy alone and with other T2R agonists in HNSCC (15,78).

### Clinical Implications of TAS2R5 Expression

The positive association between high *TAS2R5* expression and improved survival outcomes in HNSCC patients positions *TAS2R5* as a promising prognostic biomarker. Its potential utility in stratifying patients based on tumor biology could aid in personalizing treatment regimens. For instance, the expression levels of *TAS2R5* may serve as a biomarker for determining which patients require more intensive standard treatment regimens or those who can be safely de-escalated to minimize treatment side-effects. Alternatively, patients with high *TAS2R5* expression might benefit from therapies targeting this specific receptor.

Interestingly, *TAS2R5* expression did not differ between HPV+ and HPV− tumors, suggesting that its therapeutic relevance may extend across HNSCC subtypes. This is particularly significant for HPV− HNSCC, which is associated with poorer prognoses (79). The involvement of T2R5 in immune cell populations, such as macrophages (80–82) raises additional questions about its role in the tumor microenvironment. Understanding whether T2R5 contributes to tumor immune cell infiltration or function could provide further insights into its dual role as a therapeutic target and a biomarker (83).

### Limitations and Future Directions

Although this study establishes the role of T2R5 in mediating Ca^2+^ signaling, cytotoxicity, and apoptosis in HNSCC, the lack of *in vivo* models represents a key limitation. Mouse models, with their limited T2R homology to humans especially for human T2R5, pose challenges for preclinical testing (28). Future studies should prioritize xenograft models to bridge this translational gap. Additionally, the variable expression of *TAS2R5* across individuals underscores the importance of incorporating biomarker-driven approaches into prognostic or therapeutic development. Exploring the combination of T2R5-targeted ligands with other treatments, such as immune checkpoint inhibitors, could further expand the therapeutic landscape for HNSCC.

## Conclusion

This study identifies T2R5 as a novel therapeutic target in HNSCC. Phendione is a potent T2R5 activator capable of inducing sustained Ca^2+^ responses, reducing cell viability, and promoting apoptosis. The association between high *TAS2R5* expression and improved survival demonstrates potential for *TAS2R5* as a prognostic marker.

These findings pave the way for further exploration of T2R5 in both basic and translational cancer research, with potential to improve outcomes for HNSCC patients through precision medicine approaches.

## Acknowledgments

We thank M. Victoria (University of Pennsylvania) for excellent technical assistance and helpful discussions. This study was supported by a McCabe Foundation Fellowship Grant (RMC) and US National Institutes of Health grants R01AI167971 (RJL).

## References

1. Johnson DE, Burtness B, Leemans CR, Lui VWY, Bauman JE, Grandis JR, et al. Head and neck squamous cell carcinoma. Nature Reviews Disease Primers 2020 6:1 2020–11-26;6

2. Barsouk A, Aluru JS, PrashanthRawla, Saginala K, Barsouk A. Epidemiology, Risk Factors, and Prevention of Head and Neck Squamous Cell Carcinoma-PubMed. Medical sciences (Basel, Switzerland) 06/13/2023;11

3. Diana G, Corica C. Human Papilloma Virus vaccine and prevention of head and neck cancer, what is the current evidence? Oral Oncology 2021/04/01;115

4. Khariwala SS, Hatsukami DK, Stepanov I, Rubin N, Nelson HH. Patterns of Tobacco Cessation Attempts and Symptoms Experienced Among Smokers With Head and Neck Squamous Cell Carcinoma. JAMA Otolaryngology--Head & Neck Surgery 2018 Apr 12;144

5. Sung H, Ferlay J, Siegel RL, Laversanne M, Soerjomataram I, Jemal A, et al. Global Cancer Statistics 2020: GLOBOCAN Estimates of Incidence and Mortality Worldwide for 36 Cancers in 185 Countries. CA: a cancer journal for clinicians 2021;71:209-49

6. Bejenaru PL, Popescu B, Oancea ALA, Simion-Antonie CB, Berteșteanu GS, Condeescu-Cojocarița M, et al. Quality-of-Life Assessment after Head and Neck Oncological Surgery for Advanced-Stage Tumours. Journal of Clinical Medicine 2022 Aug 19;11

7. Sainz E, Cavenagh Margaret M, Gutierrez J, Battey James F, Northup John K, Sullivan Susan L. Functional characterization of human bitter taste receptors. Biochemical Journal 2007 Apr 12;403

8. Lang T, Pizio AD, Risso D, Drayna D, Behrens M. Activation Profile of TAS2R2, the 26th Human Bitter Taste Receptor - PubMed. Molecular nutrition & food research 2023 Jun;67

9. Lu P, Zhang C-H, Lifshitz LM, ZhuGe R. Extraoral bitter taste receptors in health and disease. The Journal of General Physiology 2017 Feb;149

10. Tuzim K, Korolczuk A, Tuzim K, Korolczuk A. An update on extra-oral bitter taste receptors. Journal of Translational Medicine 2021 19:1 2021–10-21;19

11. Jeruzal-Świątecka J, Fendler W, Pietruszewska W. Clinical Role of Extraoral Bitter Taste Receptors - PubMed. International journal of molecular sciences 07/21/2020;21

12. Zehentner S, Reiner AT, Grimm C, Somoza V, Zehentner S, Reiner AT, et al. The Role of Bitter Taste Receptors in Cancer: A Systematic Review. Cancers 2021, Vol 13, Page 5891 2021-11-23;13

13. Carey RM, McMahon D, Miller ZA, Kim T, Rajasekaran K, Gopallawa I, et al. T2R bitter taste receptors regulate apoptosis and may be associated with survival in head and neck squamous cell carcinoma - PubMed. Molecular oncology 2022 Apr;16

14. Miller ZA, Mueller A, Kim T, Jolivert J, Ma RZ, Muthuswami S, et al. Lidocaine induces apoptosis in head and neck squamous cell carcinoma through activation of bitter taste receptor T2R14 - PubMed. Cell reports 12/26/2023;42

15. Miller ZA, Muthuswami S, Mueller A, Ma RZ, Sywanycz SM, Naik A, et al. GLUT1 inhibitor BAY-876 induces apoptosis and enhances anti-cancer effects of bitter receptor agonists in head and neck squamous carcinoma cells. Cell Death Discovery 2024 Jul 25;10

16. Kim D, Woo JA, Geffken E, An SS, Liggett SB. Coupling of Airway Smooth Muscle Bitter Taste Receptors to Intracellular Signaling and Relaxation Is via Gαi1,2,3. American Journal of Respiratory Cell and Molecular Biology 2017-06-01;56

17. Ozeck M, Brust P, Xu H, Servant G. Receptors for bitter, sweet and umami taste couple to inhibitory G protein signaling pathways - PubMed. European journal of pharmacology 04/12/2004;489

18. Royds J, Khan AH, Buggy DJ. An Update on Existing Ongoing Prospective Trials Evaluating the Effect of Anesthetic and Analgesic Techniques During Primary Cancer Surgery on Cancer Recurrence or Metastasis. Int Anesthesiol Clin 2016;54:e76–83

19. Sessler DI, Pei L, Huang Y, Fleischmann E, Marhofer P, Kurz A, et al. Recurrence of breast cancer after regional or general anaesthesia: a randomised controlled trial. Lancet 2019;394:1807–15

20. Cakmakkaya OS, Kolodzie K, Apfel CC, Pace NL. Anaesthetic techniques for risk of malignant tumour recurrence. Cochrane Database Syst Rev 2014:Cd008877

21. Badwe RA, Parmar V, Nair N, Joshi S, Hawaldar R, Pawar S, et al. Effect of Peritumoral Infiltration of Local Anesthetic Before Surgery on Survival in Early Breast Cancer. J Clin Oncol 2023;41:3318–28

22. Carey R. Intratumoral Lidocaine Injection Before Oropharyngeal Cancer Surgery (856397). Clinical Trials; 2025.

23. Dagan-Wiener A, Pizio AD, Nissim I, Bahia M, Dubovski N, Margulis E, et al. BitterDB: taste ligands and receptors database in 2019 - PubMed. Nucleic acids research 01/08/2019;47

24. Pizio AD, Niv MY. Promiscuity and selectivity of bitter molecules and their receptors. Bioorganic & Medicinal Chemistry 2015/07/15;23

25. Soares S, Kohl S, Thalmann S, Mateus N, Meyerhof W, Freitas VD. Different phenolic compounds activate distinct human bitter taste receptors - PubMed. Journal of agricultural and food chemistry 02/20/2013;61

26. Meyerhof W, Batram C, Kuhn C, Brockhoff A, Chudoba E, Bufe B, et al. The Molecular Receptive Ranges of Human TAS2R Bitter Taste Receptors. Chemical Senses 2010/02/01;35

27. Devillier P, Naline E, Grassin-Delyle S. The pharmacology of bitter taste receptors and their role in human airways. Pharmacology & Therapeutics 2015/11/01;155

28. Grau-Bové C, Grau-Bové X, Terra X, Garcia-Vallve S, Rodríguez-Gallego E, Beltran-Debón R, et al. Functional and genomic comparative study of the bitter taste receptor family TAS2R: Insight into the role of human TAS2R5. The FASEB Journal 2022/03/01;36

29. Kim D, An S, Lam H, Leahy J, Liggett S. Identification and Characterization of Novel Bronchodilator Agonists Acting at Human Airway Smooth Muscle Cell TAS2R5 - PubMed. ACS pharmacology & translational science 11/05/2020;3

30. Viganor L, Galdino ACM, Nunes APF, Santos KRN, Branquinha MH, Devereux M, et al. Anti-Pseudomonas aeruginosa activity of 1,10-phenanthroline-based drugs against both planktonic-and biofilm-growing cells. Journal of Antimicrobial Chemotherapy 2016/01/01;71

31. McCann M, Kellett A, Kavanagh K, Devereux M, Santos A. Deciphering the antimicrobial activity of phenanthroline chelators - PubMed. Current medicinal chemistry 2012;19

32. Meade E, Savage M, Slattery M, Garvey M. Investigation of Alternative Therapeutic and Biocidal Options to Combat Antifungal-Resistant Zoonotic Fungal Pathogens Isolated from Companion Animals. Infectious Disease Reports 2021 Apr 11;13

33. Granato M, Gonçalves D, Seabra S, McCann M, Devereux M, Santos AD, et al. 1,10-Phenanthroline-5,6-Dione-Based Compounds Are Effective in Disturbing Crucial Physiological Events of Phialophora verrucosa - PubMed. Frontiers in microbiology 01/30/2017;8

34. Deegan C, Coyle B, McCann M, Devereux M, Egan DA. In vitro anti-tumour effect of 1,10-phenanthroline-5,6-dione (phendione), [Cu(phendione)3](ClO4)2·4H2O and [Ag(phendione)2]ClO4 using human epithelial cell lines. Chemico-Biological Interactions 2006/12/01;164

35. Yue J, Vendramin R, Liu F, Lopez O, Valencia M, Santos HGD, et al. Targeted chemotherapy overcomes drug resistance in melanoma-PubMed. Genes & development 05/01/2020;34

36. Sears CR, Turchi JJ. Complex Cisplatin-Double Strand Break (DSB) Lesions Directly Impair Cellular Non-Homologous End-Joining (NHEJ) Independent of Downstream Damage Response (DDR) Pathways. The Journal of Biological Chemistry 2012 May 23;287

37. Farhangian H, Kharat AN. Novel ternary Pd(II) complexes as potent anticancer drugs for ovarian carcinoma treatment. Journal of Molecular Structure 2023/12/15;1294

38. Schindelin J, Arganda-Carreras I, Frise E, Kaynig V, Longair M, Pietzsch T, et al. Fiji: an open-source platform for biological-image analysis. Nature Methods 2012 9:7 2012-06-28;9

39. Christensen J, Schmidt H, Steiniche T, Madsen M. Identification of robust reference genes for studies of gene expression in FFPE melanoma samples and melanoma cell lines - PubMed. Melanoma research 2020 Feb;30

40. Wu J, Prole D, Shen Y, Lin Z, Gnanasekaran A, Liu Y, et al. Red fluorescent genetically encoded Ca2+ indicators for use in mitochondria and endoplasmic reticulum - PubMed. The Biochemical journal 11/15/2014;464

41. Kenerson HL, Sullivan KM, Seo YD, Stadeli KM, Ussakli C, Yan X, et al. Tumor slice culture as a biologic surrogate of human cancer. Annals of Translational Medicine 2020 Feb;8

42. Park J, Kim K-S, Kim K-H, Lee I-S, Jeong H-s, Kim Y, et al. GLP-1 secretion is stimulated by 1,10-phenanthroline via colocalized T2R5 signal transduction in human enteroendocrine L cell. Biochemical and Biophysical Research Communications 2015/12/04;468

43. Delpapa EH, Simas TM, ZhuGe R, Qu M. Bitter taste receptor TAS2R5 agonist phenanthroline as a broad-spectrum uterine relaxant. American Journal of Obstetrics & Gynecology 2023/01/01;228

44. Talmon M, Pollastro F, Fresu LG. The Complex Journey of the Calcium Regulation Downstream of TAS2R Activation. Cells 2022 Nov 16;11

45. Workman AD, Palmer JN, Adappa ND, Cohen NA. The Role of Bitter and Sweet Taste Receptors in Upper Airway Immunity. Current allergy and asthma reports 2015 Dec;15

46. Soares S, Brandão E, Guerreiro C, Soares S, Mateus N, Freitas Vd. Tannins in Food: Insights into the Molecular Perception of Astringency and Bitter Taste. Molecules 2020 Jun 2;25

47. Yang MY, Kim S-K, Kim D, Liggett SB, William A Goddard I. Structures and Agonist Binding Sites of Bitter Taste Receptor TAS2R5 Complexed with Gi Protein and Validated against Experiment. The journal of physical chemistry letters 2021 Sep 20;12

48. Soares S, Silva MS, García-Estevez I, Groβmann P, Brás N, Brandão E, et al. Human Bitter Taste Receptors Are Activated by Different Classes of Polyphenols. Journal of Agricultural and Food Chemistry July 28, 2018;66

49. Uemura T, Kawasaki T, Taniguchi M, Moritani Y, Hayashi K, Saito T, et al. Biological properties of a specific Gαq/11 inhibitor, YM-254890, on platelet functions and thrombus formation under high-shear stress. British Journal of Pharmacology 2006 Mar 6;148

50. Schrage R, Schmitz A, Gaffal E, Annala S, Kehraus S, Wenzel D, et al. The experimental power of FR900359 to study Gq-regulated biological processes. Nature Communications 2015 6:1 2015-12-14;6

51. Fierro F, Peri L, Hübner H, Tabor-Schkade A, Waterloo L, Löber S, et al. Inhibiting a promiscuous GPCR: iterative discovery of bitter taste receptor ligands. Cellular and Molecular Life Sciences: CMLS 2023 Apr 3;80

52. Odaka K, Arai S, Inoue T, Kitaguchi T. Genetically-encoded yellow fluorescent cAMP indicator with an expanded dynamic range for dual-color imaging - PubMed. PloS one 06/24/2014;9

53. McMahon DB, Kuek LE, Johnson ME, Johnson PO, Horn RLJ, Carey RM, et al. The bitter end: T2R bitter receptor agonists elevate nuclear calcium and induce apoptosis in non-ciliated airway epithelial cells. Cell Calcium 2022/01/01;101

54. Freund J, Mansfield C, Doghramji L, Adappa N, Palmer J, Kennedy D, et al. Activation of airway epithelial bitter taste receptors by Pseudomonas aeruginosa quinolones modulates calcium, cyclic-AMP, and nitric oxide signaling - PubMed. The Journal of biological chemistry 06/22/2018;293

55. Cui H, Green RD. Regulation of the cAMP-elevating effects of isoproterenol and forskolin in cardiac myocytes by treatments that cause increases in cAMP. Biochemical and Biophysical Research Communications 2003/07/18;307

56. Vardjan N, Kreft M, Zorec R. Dynamics of β-adrenergic/cAMP signaling and morphological changes in cultured astrocytes - PubMed. Glia 2014 Apr;62

57. Deegan C, Coyle B, McCann M, Devereux M, Egan D. In vitro anti-tumour effect of 1,10-phenanthroline-5,6-dione (phendione), [Cu(phendione)3](ClO4)2.4H2O and [Ag(phendione)2]ClO4 using human epithelial cell lines - PubMed. Chemico-biological interactions 12/01/2006;164

58. Yang MY, Mac KD, Strzelinski HR, Hoffman SA, Kim D, Kim S-K, et al. Agonist activation to open the Gα subunit of the GPCR–G protein precoupled complex defines functional agonist activation of TAS2R5. Proceedings of the National Academy of Sciences 2024-11-20;121

59. Harris JC, Lee RJ, Carey RM, Harris JC, Lee RJ, Carey RM. Extragustatory bitter taste receptors in head and neck health and disease. Journal of Molecular Medicine 2024 102:12 2024-09-25;102

60. D’Urso O, Drago F. Pharmacological significance of extra-oral taste receptors. European Journal of Pharmacology 2021/11/05;910

61. Welcome MO, Mastorakis NE. The taste of neuroinflammation: Molecular mechanisms linking taste sensing to neuroinflammatory responses. Pharmacological Research 2021/05/01;167

62. Shaw L, Mansfield C, Colquitt L, Lin C, Ferreira J, Emmetsberger J, et al. Personalized expression of bitter’taste’ receptors in human skin - PubMed. PloS one 10/17/2018;13

63. Hajnóczky G, Csordás G, Das S, Garcia-Perez C, Saotome M, Roy SS, et al. Mitochondrial calcium signalling and cell death: approaches for assessing the role of mitochondrial Ca2+ uptake in apoptosis. Cell calcium 2006 Oct 30;40

64. Görlach A, Bertram K, Hudecova S, Krizanova O. Calcium and ROS: A mutual interplay. Redox Biology 2015 Aug 11;6

65. Hempel N, Trebak M. Crosstalk between Calcium and Reactive Oxygen Species Signaling in Cancer. Cell calcium 2017 Jan 18;63

66. Pinton P, Giorgi C, Siviero R, Zecchini E, Rizzuto R. Calcium and apoptosis: ER-mitochondria Ca2+ transfer in the control of apoptosis. Oncogene 2008 Oct 27;27

67. Lee Y. Cancer Chemopreventive Potential of Procyanidin. Toxicological Research 2015 Oct 15;33

68. Prasad R, Katiyar SK. Bioactive Phytochemical Proanthocyanidins Inhibit Growth of Head and Neck Squamous Cell Carcinoma Cells by Targeting Multiple Signaling Molecules. PLOS ONE Sep 26, 2012;7

69. Nandakumar V, Singh T, Katiyar SK. Multi-targeted prevention and therapy of cancer by proanthocyanidins. Cancer letters 2008 May 23;269

70. Sun L, Subar AF, Bosire C, Dawsey SM, Kahle LL, Zimmerman TP, et al. Dietary Flavonoid Intake Reduces the Risk of Head and Neck but Not Esophageal or Gastric Cancer in US Men and Women. The Journal of Nutrition 2017 Jul 19;147

71. Sriram K, Insel PA. G Protein-Coupled Receptors as Targets for Approved Drugs: How Many Targets and How Many Drugs? Molecular Pharmacology 2018 Apr;93

72. Peri L, Matzov D, Huxley DR, Rainish A, Fierro F, Sapir L, et al. A bitter anti-inflammatory drug binds at two distinct sites of a human bitter taste GPCR. Nature Communications 2024 Nov 18;15

73. Hanahan D, Weinberg Robert A. Hallmarks of Cancer: The Next Generation. Cell 2011/03/04;144

74. Raudenská M, Balvan J, Masařík M, Raudenská M, Balvan J, Masařík M. Cell death in head and neck cancer pathogenesis and treatment. Cell Death & Disease 2021 12:2 2021-02-18;12

75. Chaudhry G, Akim A, Sung Y, Muhammad T. Cancer and Apoptosis - PubMed. Methods in molecular biology (Clifton, NJ) 2022;2543

76. Molinolo AA, Amornphimoltham P, Squarize CH, Castilho RM, Patel V, Gutkind JS. Dysregulated Molecular Networks in Head and Neck Carcinogenesis. Oral oncology 2008 Sep 19;45

77. Neophytou CM, Trougakos IP, Erin N, Papageorgis P. Apoptosis Deregulation and the Development of Cancer Multi-Drug Resistance. Cancers 2021 Aug 28;13

78. Mazambani S, Choe JH, Oh T-G, Singh PK, Kim J-w, Kim TH. Targeting the glucose-insulin link in head and neck squamous cell carcinoma induces cytotoxic oxidative stress and inhibits cancer growth. bioRxiv 2023-07-15

79. Sabatini ME, Chiocca S, Sabatini ME, Chiocca S. Human papillomavirus as a driver of head and neck cancers. British Journal of Cancer 2019 122:3 2019–11- 11;122

80. Harmon CP, Deng D, Breslin PA. Bitter Taste Receptors (T2Rs) are Sentinels that Coordinate Metabolic and Immunological Defense Responses. Current opinion in physiology 2021 Jan 12;20

81. Grassin-Delyle S, Salvator H, Mantov N, Abrial C, Brollo M, Faisy C, et al. Bitter Taste Receptors (TAS2Rs) in Human Lung Macrophages: Receptor Expression and Inhibitory Effects of TAS2R Agonists - PubMed. Frontiers in physiology 10/02/2019;10

82. Malki A, Fiedler J, Fricke K, Ballweg I, Pfaffl M, Krautwurst D. Class I odorant receptors, TAS1R and TAS2R taste receptors, are markers for subpopulations of circulating leukocytes - PubMed. Journal of leukocyte biology 2015 Mar;97

83. Gopallawa I, Freund JR, Lee RJ. Bitter taste receptors stimulate phagocytosis in human macrophages through calcium, nitric oxide, and cyclic-GMP signaling. Cellular and Molecular Life Sciences: CMLS 2020 Mar 14;78

