## Supplemental Figures for "T2R5 agonist phendione decreases cell viability and induces apoptosis in head and neck squamous cell carcinoma"

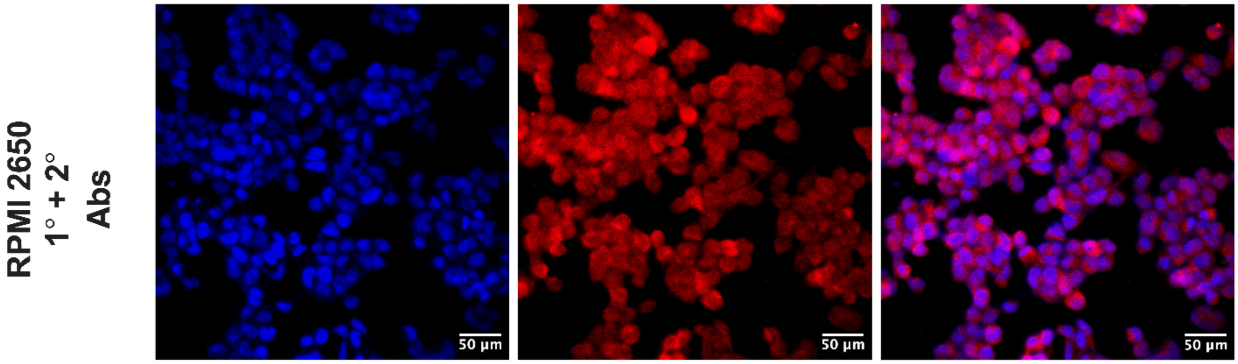


**Supplemental Figure 1: T2R5 expression in RPMI2650.**

T2R5 protein expression in RPMI2650 measured via immunofluorescence in SCC47 (DAPI = nucleus).

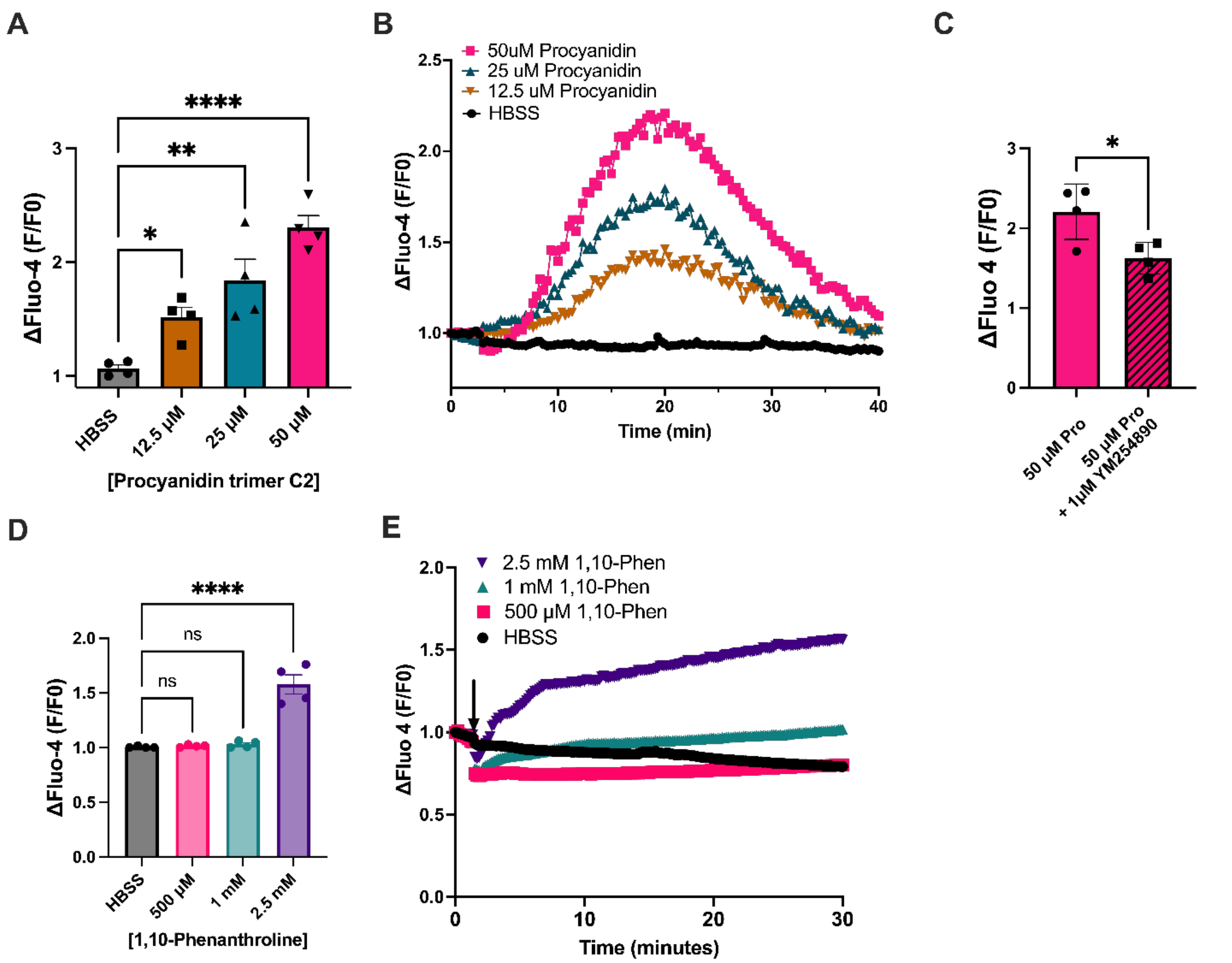


**Supplemental Figure 2: $Ca^{2+}$ responses with known T2R5-specific agonists.**

**A-B** SCC47 were incubated with Fluo-4 and stimulated with procyanidin trimer C2 leading to a significant $Ca^{2+}$ response that was dose dependent. Significance by one-way ANOVA with Dunnett's multiple comparison. **C** $Ca^{2+}$ response from procyanidin (pro) was significantly reduced with Gαq inhibitor YM254890. Significance by unpaired t-test. **D-E** Stimulation with parent compound, 1,10-phenanthroline (1,10-phen) results in a significant dose-dependent $Ca^{2+}$ response, although at much greater concentrations than phendione. *p < 0.05, **p < 0.01, ****p < 0.0001; ns, no statistical significance.

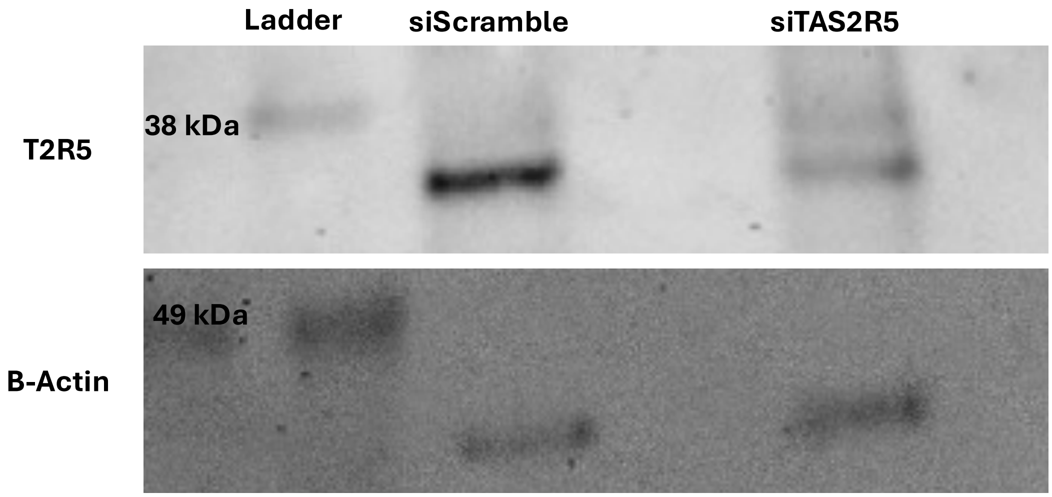


**Supplemental Figure 3: T2R5 siRNA confirmation.**

T2R5 protein knockdown in siTAS2R5 versus control (siScramble, non-targeting human).

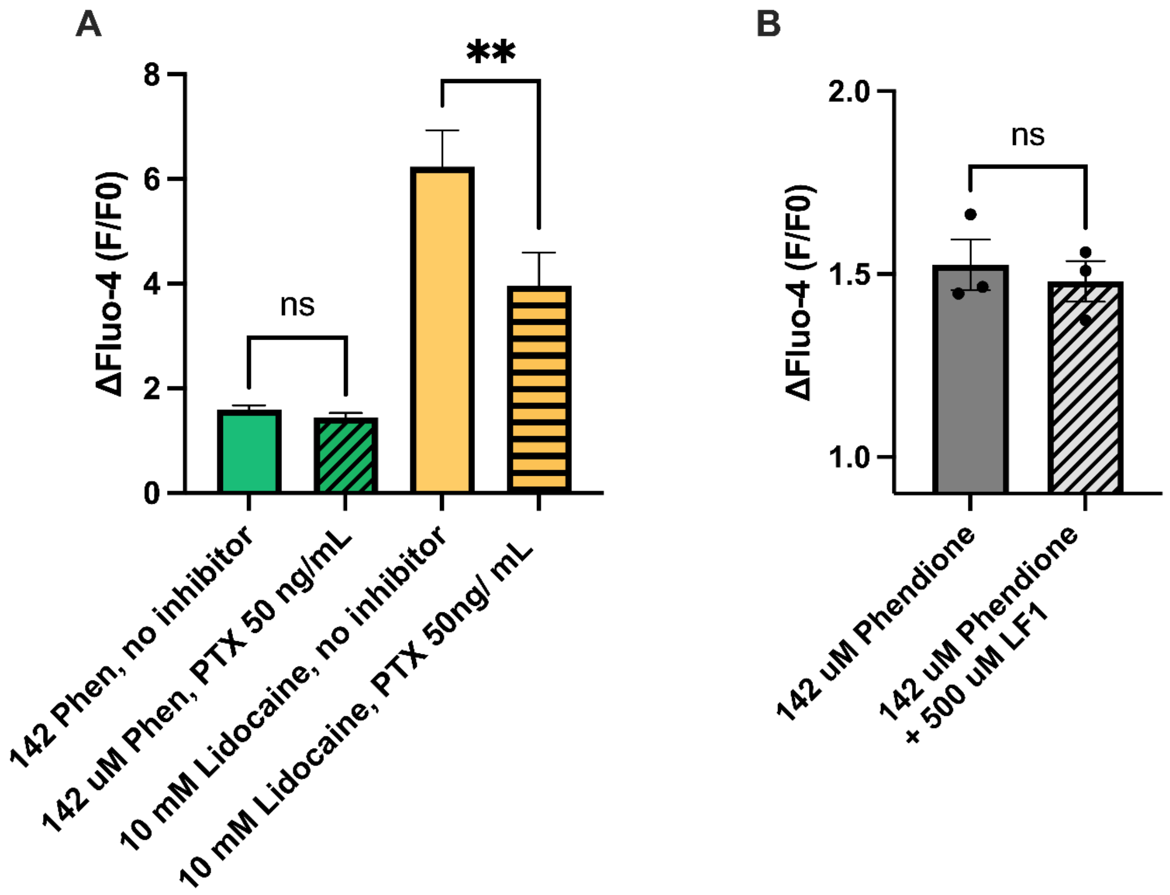


**Supplemental Figure 4: Phendione acts on a pathway distinct from T2R14.**

**A** Cells were incubated with and without $G\alpha_{i/o}$ inhibitor, pertussis toxin (PTX) at 50 ng/mL. Unlike T2R14 agonist lidocaine, there was no significant difference in $Ca^{2+}$ response with phendione stimulation. Significance by t-test, n =3, mean peak calcium response ± SEM. **B** No significant reduction in peak $Ca^{2+}$ response after phendione stimulation was observed in the presence of T2R14 antagonist LF1. Bars represent mean peak calcium ± SEM.

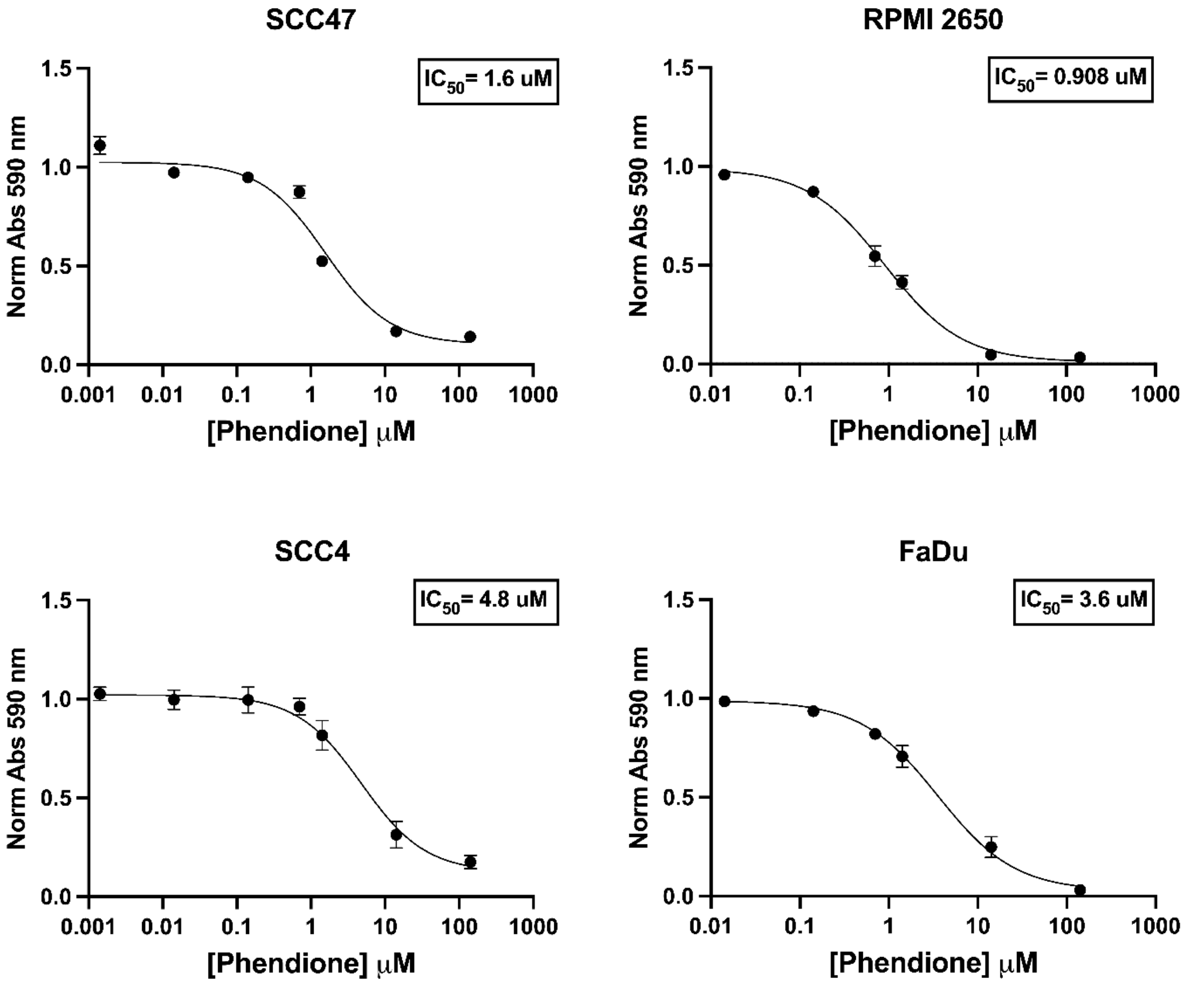


**Supplemental Figure 5: Dose dependent reduction in cell viability after 24-hour exposure to phendione.**

HNSCC cell lines have reduced cell viability in a dose dependent manner with $IC_{50}$ values ranging from 0.8-4.8 µM. IC50 values calculated using Prism software.

| Age | Sex | Primary Site | p16 |
|---|---|---|---|
| 72.4 | Male | Tonsil (palatine) | + |
| 51.1 | Male | Lateral tongue | + |
| 66.6 | Male | Tonsil (palatine) | + |
| 77.0 | Male | Tonsil (palatine) | + |
| 55.4 | Male | Buccal mucosa | - |

**Supplemental Table 1: Demographic table of HNSCC tumor slices used for cell viability assay (Figure 4H).**

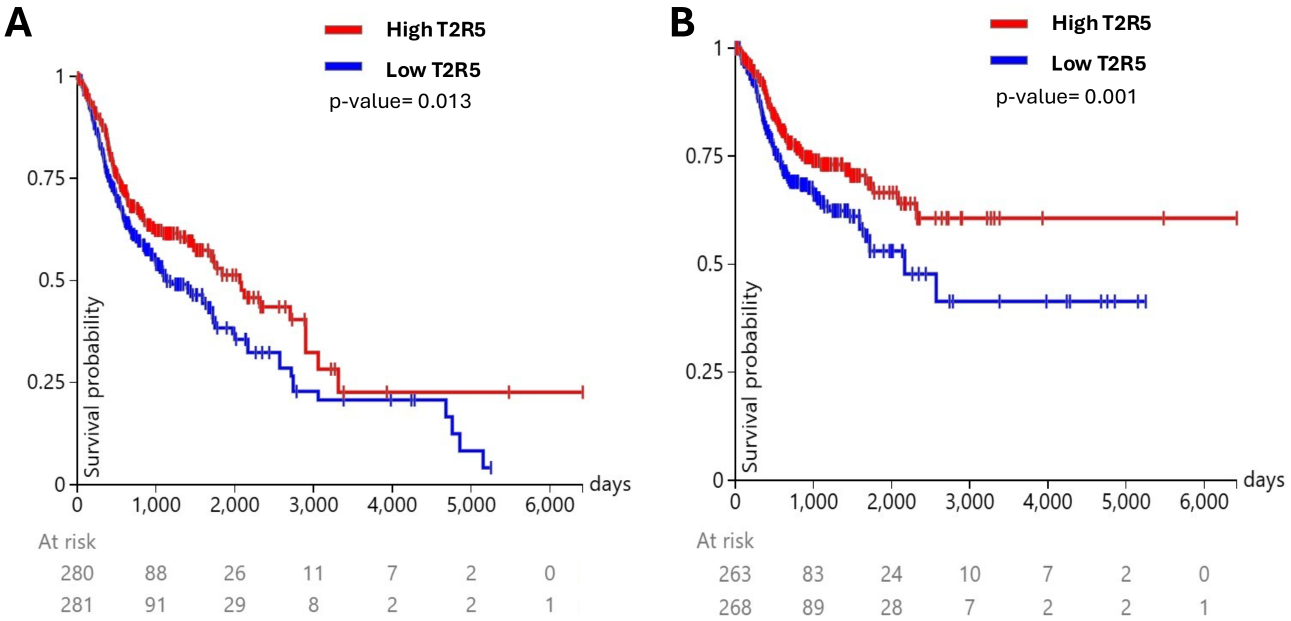


**Supplemental Figure 6:** Using bulk RNA-sequencing data from The Cancer Genome Atlas (TCGA), HNSCC patients were divided into groups based on high and low *TAS2R5* expression. Kaplan–Meier survival analysis demonstrated improved 10-year overall survival (A) and disease-specific survival (B) for cases with high *TAS2R5* expression compared to low *TAS2R5* expression (p = 0.013 and 0.001 by log-rank test, respectively).
